# The RNA helicase Ded1 regulates translation and granule formation during multiple phases of cellular stress responses

**DOI:** 10.1101/2021.06.02.446858

**Authors:** Peyman P. Aryanpur, Telsa M. Mittelmeier, Timothy A. Bolger

## Abstract

Ded1 is a conserved RNA helicase that promotes translation initiation in steady-state conditions. Ded1 has also been shown to regulate translation during cellular stress and affect the dynamics of stress granules (SGs), accumulations of RNA and protein linked to translation repression. To better understand its role in stress responses, we examined Ded1 function in two different models: *DED1* overexpression and oxidative stress. *DED1* overexpression inhibits growth and promotes the formation of SGs. A *ded1*mutant lacking the low-complexity C-terminal region (*ded1-ΔCT*), which mediates Ded1 oligomerization and interaction with the translation factor eIF4G, suppressed these phenotypes, consistent with other stresses. During oxidative stress, a *ded1-ΔCT* mutant was defective in growth and in SG formation compared to wild-type cells, although SGs were increased rather than decreased in these conditions. Unlike stress induced by direct TOR inhibition, the phenotypes in both models were only partially dependent on eIF4G interaction, suggesting an additional contribution from Ded1 oligomerization. Furthermore, examination of the growth defects and translational changes during oxidative stress suggested that Ded1 plays a role during recovery from stress. Integrating these disparate results, we propose that Ded1 controls multiple aspects of translation and RNP dynamics in both initial stress responses and during recovery.

## Introduction

Organisms are frequently subjected to various adverse conditions, including a lack of nutrients, chemical imbalances, and exposure to toxic substances. In order to survive and adapt to these stresses, cells employ a variety of responses, including highly coordinated changes in gene expression (1–3). Changes in translation play a particularly large role in stress responses because of their ability to rapidly alter the proteome and the need to reduce the outsized energy requirements of protein synthesis during stress. A number of different signaling mechanisms converge on the translation machinery in stress conditions, including activation of the integrated stress response (ISR) that results in inhibitory phosphorylation of the translation factor eIF2 and down-regulation of the target-of-rapamycin (TOR) pathway that integrates growth signals in a variety of conditions (4, 5). Translation of most mRNAs, especially those associated with growth, is greatly diminished during stress conditions (6). On the other hand, “stress-responsive” genes are up-regulated, and ribosome profiling of stressed cells has shown increases in occupancy at upstream open reading frames (uORFs) and non-AUG initiation (7, 8). These data suggest that the translational stress response is complex and specific, but the sources of this specificity are not fully clear.

A common feature of many types of stress conditions is the formation of stress granules (SGs), non-membranous organelles composed of RNAs and proteins (9). SGs appear to form as a result of multiple interactions amongst these components, often including proteins with low-complexity, intrinsically-disordered regions (IDRs) that promote liquid-liquid phase separation (10). Despite close association with both stress conditions and translation repression, the function of SGs is not well understood, although mutations in genes required for SG formation also reduce cell survival in stress (9). Stress granules are often suggested to function as sorting and storage sites for mRNAs during stress (11). Consistent with this hypothesis, RNA-seq and localization studies have shown that some mRNAs are enriched in SGs while others are depleted, and mRNAs are able to actively exchange between SGs, the cytosol, and other structures (12–14). The mRNAs in SGs have generally been considered to be translationally-repressed, although a recent study has suggested that this may not always be the case (15).

Ded1 is a translation factor in budding yeast that plays several different roles in translation initiation (16, 17). Ded1 (DDX3X in humans) is a member of the DEAD-box RNA helicase family, which utilize ATP to alter RNA-RNA and RNA-protein interactions and are critical for many gene expression processes (18). Similar to many DEAD-box proteins, the Ded1 domain structure consists of a central helicase core flanked by long extensions that are predicted to be IDRs (Figure 1A). These N- and C-terminal regions mediate binding to other proteins, including members of the eIF4F translation complex, and oligomerization of Ded1 itself (19–23). Canonically, Ded1 stimulates translation initiation in steady-state growth conditions by unwinding secondary structure in 5’ UTRs, facilitating start site scanning by the translation pre-initiation complex (PIC). Thus, mRNAs with more complex, structured 5’ UTRs tend to be hyper-dependent on Ded1 activity, while those with simpler 5’ UTRs are less affected (24, 25). Furthermore, *ded1* mutation or depletion results in utilization of alternative translation initiation sites (ATIS) that may affect downstream translation or protein function (25). Ded1 also promotes assembly of the 43S PIC on mRNA, again in an mRNA-specific manner (19, 23, 26, 27).

**Figure 1:**
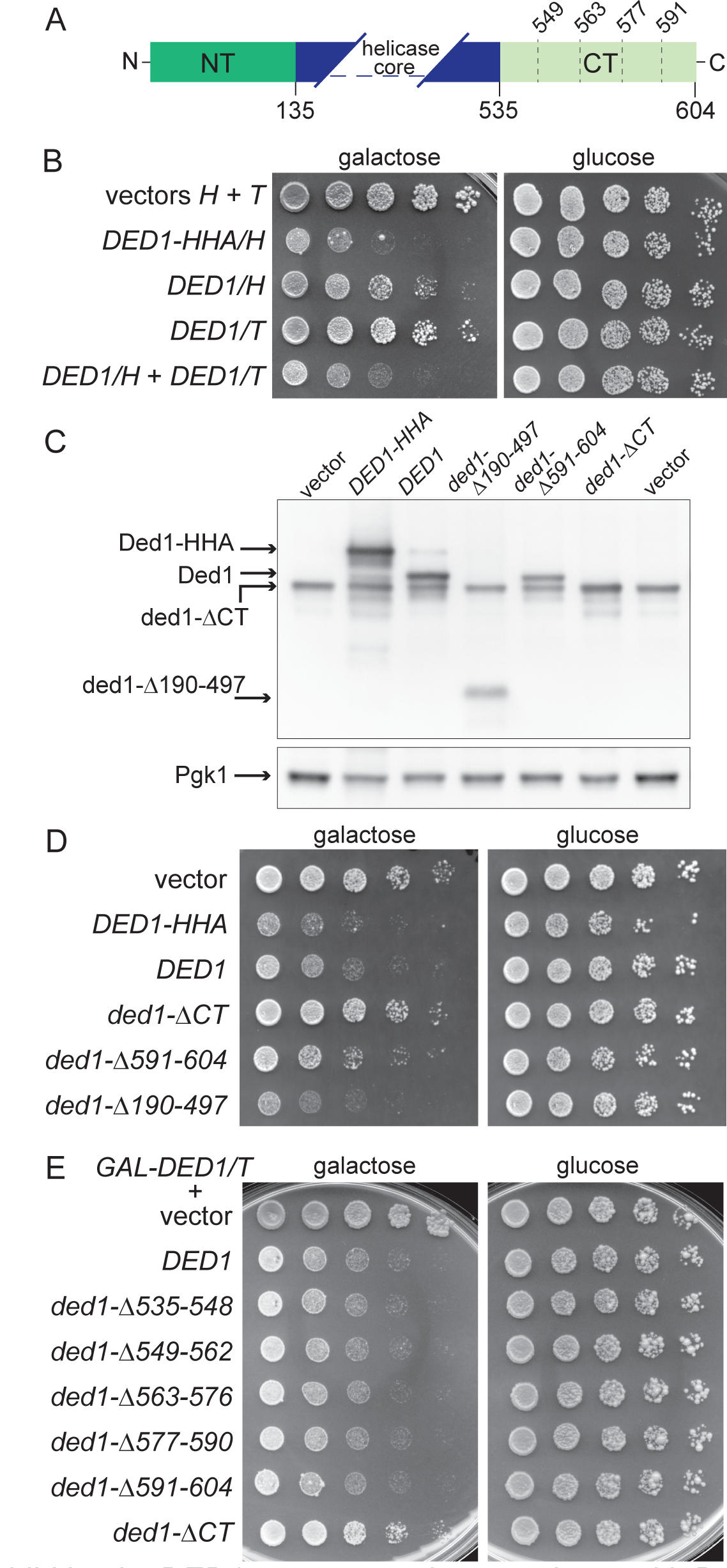
Growth inhibition by DED1 overexpression correlates with Ded1 protein levels and requires the C-terminal region. (A) Domain map of the Ded1 protein illustrating the N- and C-terminal low-complexity regions (NT and CT, respectively) and the boundaries of five sequential deletions of 14 residues within the CT analyzed in this study. (B) Five-fold serial dilutions of *ded1-ΔCT* cells with galactose-inducible wild-type *DED1* spotted on selective medium containing either galactose or glucose. Cells harbored a single plasmid that encoded wild-type Ded1 with a C-terminal tag (*DED1-HHA*), a single plasmid encoding untagged Ded1 (*DED1/H* or *DED1/T*), or two plasmids that each encoded untagged Ded1 (*DED1/H + DED1/T*). (C) Western blot analysis of protein extracts from *ded1-ΔCT* cells containing the indicated plasmid, induced for 7 hours. Samples were probed with antibodies specific for Ded1 or Pgk1 (as a loading control). (D) Five-fold serial dilutions of *ded1-ΔCT* cells with galactose-inducible *DED1* or *ded1* mutants spotted on selective medium containing either galactose or glucose. (E) Five-fold serial dilutions of *ded1-ΔCT* cells containing galactose-inducible *DED1* (*DED1/T*) plus a second galactose-inducible construct as indicated, spotted on selective medium containing either galactose or glucose.

Ded1 and its orthologs have also been implicated in translation repression during stress conditions and are associated with SGs in particular. Ded1 and DDX3X are major protein components of SGs and can affect SG assembly (19, 28, 29). Overexpression of *DED1* alone can induce SG-like foci in cells (19), while Ded1 undergoes phase separation *in vitro* under conditions of high Ded1 concentration, elevated temperature, and/or the presence of RNA (30, 31). These effects are at least partially mediated by the N- and C-terminal IDRs (19, 30, 31). Overall, Ded1 appears to promote SG formation, although the consequences of this stimulation have not been fully defined. Iserman et al. have proposed that sequestration of Ded1 into granules during heat shock causes a switch in translation to mRNAs with less complex 5’ UTRs, but this model has not been fully tested (31).

Recently, we showed that Ded1 has a role downstream of the TOR pathway in the translational stress response (32). Specifically, when the TOR pathway was down-regulated, Ded1 was required for efficient translation repression and growth inhibition. In particular, the low-complexity C-terminal region of Ded1 was necessary for these effects, which promoted degradation of its binding partner eIF4G. We proposed that in these conditions, Ded1 remodeled translation complexes to cause dissociation and degradation of eIF4G, thus reducing bulk translation during stress. However, it is currently unclear how Ded1’s role in SG dynamics may affect this model, particularly in different cellular stresses. Here we sought to address these gaps by examining Ded1-dependent mechanisms of translation regulation in two different conditions. First, upon *DED1* overexpression, both growth inhibition and SG formation were dependent on Ded1 levels and the presence of the C-terminal region, but not helicase activity. Furthermore, Ded1 interaction with eIF4G had only a modest effect, suggesting a contribution from Ded1 oligomerization. Second, when cells were subjected to oxidative stress through addition of peroxide to the media, the C-terminal region played critical roles both in survival as well as the ability of cells to recovery from the stress over time. Interestingly, deletion of the C-terminal region increased SG formation during oxidative stress, in a manner opposite to the overexpression results. Similar to overexpression, Ded1 interaction with eIF4G played a moderate role in responding to oxidative stress. Consistent with the effects on growth and SG formation, reporter assays revealed that Ded1 and its C-terminal region mediate changes in translation during a time course of peroxide treatment. To integrate these results, we propose a biphasic model for the function of Ded1 in the stress response wherein it has distinct functions on cell survival/growth, translation, and SGs in an early response phase and during a later recovery/adaptation phase.

## Results

### DED1 growth inhibition is dependent on protein levels and the C-terminal region but not the helicase domain

We recently showed that Ded1 plays a role in the translational response to TOR pathway inhibition in a manner dependent on its C-terminal domain and interaction with eIF4G (32). To examine whether this mechanism is similar during other conditions of translation repression, including SG induction, we first utilized overexpression of *DED1*. Hilliker et al. previously demonstrated that *DED1* overexpression causes growth inhibition and formation of SG-like aggregates, dependent on the presence of conserved “assembly domains” (19). Likewise, we observed severe growth inhibition upon overexpression of tagged, wild-type *DED1* from a galactose-inducible promoter (Figure 1B, *DED1-HHA*). However, when untagged *DED1* was expressed from a similar plasmid backbone, the growth inhibition, while still present, was much less severe (Figure 1B, *DED1/H*). Western blotting of the expressed constructs showed that Ded1 protein levels of the tagged version were about twice that of the untagged *DED1*, suggesting that the C-terminal tag stabilizes the protein (Figure 1C). This result further suggests that the growth inhibition is quite sensitive to Ded1 protein levels. Supporting this idea, overexpression of *DED1* from two untagged plasmids in the same cells caused a greater decrease in growth, similar to tagged *DED1* (Figure 1B, *DED1/H + DED1/T*). Untagged *DED1* was used for the remainder of this study unless otherwise noted.

We next sought to determine whether the functional requirements we identified in TOR pathway downregulation also affect *DED1*-mediated growth inhibition. Consistent with previous results (19), deletion of the C-terminal region, *ded1-ΔCT (Δ536-604)*, largely abrogated the ability of Ded1 to inhibit growth (Figure 1D). In our prior study, a smaller deletion of the final 14 amino acids of the C-terminal region (*ded1-Δ591-604*), which greatly reduced binding to eIF4G *in vitro*, phenocopied the larger deletion (32). In contrast with these results after TOR inhibition, overexpressing the *ded1-Δ591-604* mutant inhibited growth only slightly less than wild-type *DED1* (Figure 1D), indicating that there are differences between the Ded1-dependent mechanisms. Likewise, each of a set of similar 14 amino acid deletions in the C-terminal region showed growth inhibition similar to wild-type *DED1*, either in combination with another *DED1* overexpression plasmid (Figure 1E) or alone (Supplemental Figure S1). Furthermore, we constructed a mutant that deleted most of the central helicase domain, *ded1-Δ190-497*, leaving the N- and C-terminal domains fused together with only short flanking sequences (Figure 1A).

This mutant inhibited growth to a slightly greater extent than wild-type *DED1* (Figure 1D). It was previously shown that growth inhibition did not require Ded1 activity, as an ATPase-deficient mutant had a similar phenotype to wild-type *DED1* (19), and this result extends that conclusion by suggesting that the N- and C-terminal regions themselves are sufficient for these effects. Again, this differs from our prior results in TOR inhibition wherein Ded1 activity was required for the effects on translation repression (32).

### Ded1-induced SGs are dependent on the C-terminal region but not the helicase domain

Overexpression of *DED1* has been shown to induce SG-like foci that contain a number of SG components, including mRNAs and translation factors, and formation of the Ded1-induced SGs correlates with growth inhibition (19). Therefore, we tested whether SG formation was affected by the *ded1* mutations examined above using a *PAB1-GFP* reporter, a well-established yeast SG marker (33). After 7 hours of galactose induction, *DED1*-overexpressing cells frequently contained one or more Pab1-GFP-positive foci, indicating SG formation in 24% of cells, while very few SGs were observed in control cells (Figure 2A, B). Further increasing *DED1* levels (with a second overexpression plasmid) led to an additional increase in the percentage of cells with SGs, consistent with the inhibitory effects on growth. In contrast, cells expressing the *ded1-ΔCT* mutant contained almost no SGs, similar to the control cells. However, both the *ded1-Δ591-604* and *Δ190-497* mutants both displayed rates of SG induction similar to the wild-type *DED1*-expressing cells (Figure 2B). Interestingly, we noted that the Pab1-GFP foci induced by the *ded1-Δ190-497* mutant had a somewhat different qualitative appearance with individual foci more elongated and extended compared to the largely round SGs in the wild-type *DED1-*expressing cells (Supplemental Figure S2). This phenotype is also distinct from that with the tagged version of *DED1*, which were often very large but were still more rounded than the *ded1-Δ190-497* mutant. Overall, these results indicate that the Ded1 C-terminal region plays an important role in inducing SGs, while, surprisingly, the helicase domain does not. Furthermore, the SG results closely correlate with the growth inhibition shown in Figure 1, suggesting a functional relationship between growth inhibition and SG formation.

**Figure 2:**
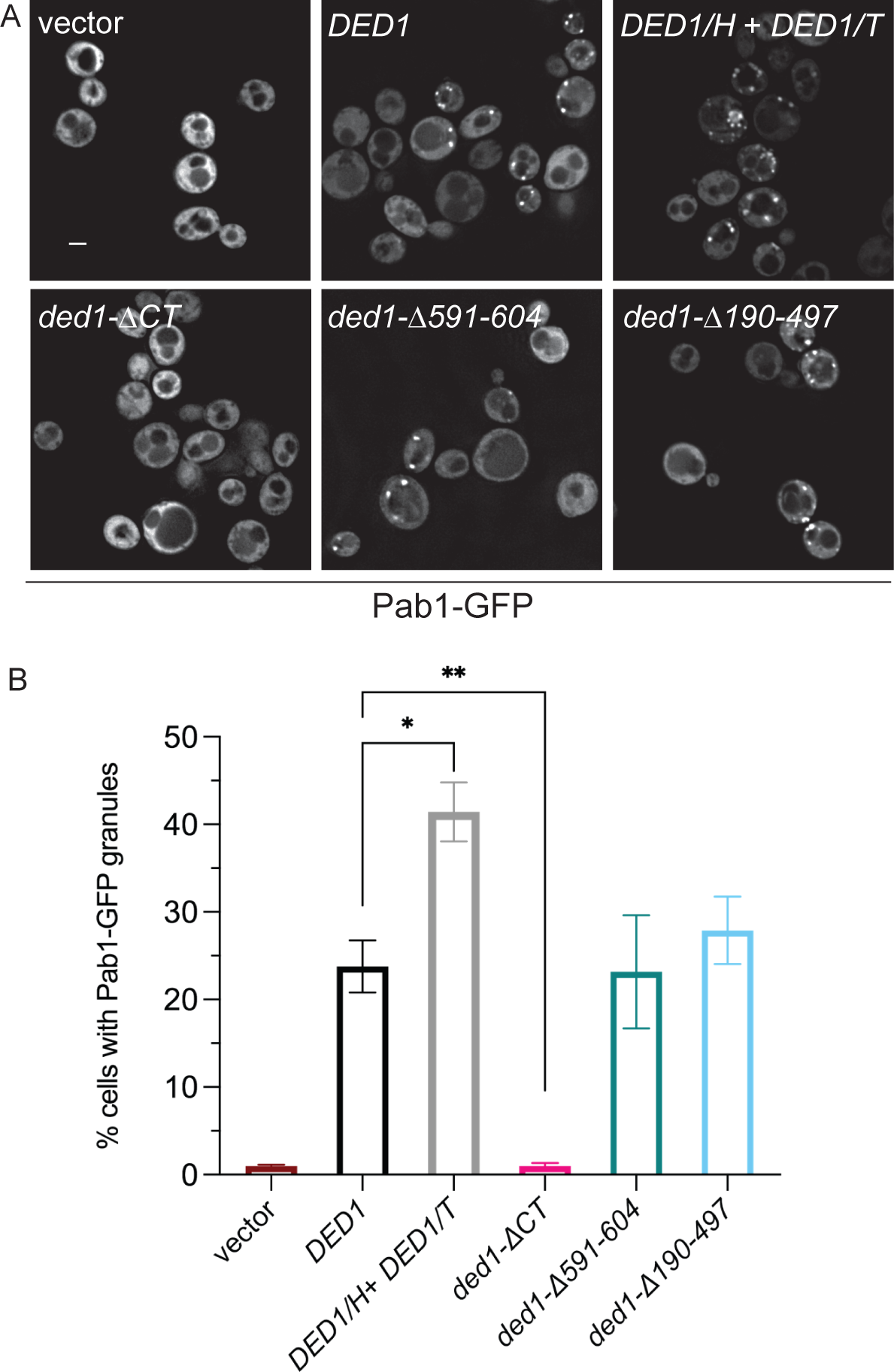
The Ded1 C-terminus is required for formation of GAL-DED1-induced granules. (A) Live-cell microscopy showing Pab1-GFP granules in *ded1-ΔCT* cells expressing galactose-inducible wild-type *DED1* from a single plasmid construct (*DED1*), two inducible constructs (*DED1/H* + *DED1/T*), or the indicated *ded1* deletion mutant constructs. Cells were grown in liquid medium containing galactose for 7 hours before imaging. Scale bar = 2 μm. (B) Quantitation of the presence of Pab1-GFP granules as the percentage of cells that contained GFP-positive foci. Mean and SEM of 3-11 replicates are shown. Statistical significance was determined using Student’s t-test (unpaired; * p < 0.05, ** p < 0.01).

### Ded1 interaction with eIF4G has moderate effects on DED1 overexpression phenotypes

The C-terminal domain of Ded1 has been shown to both interact with eIF4G and mediate Ded1 self-oligomerization, but how these individual interactions affect Ded1 function *in vivo* remain unclear (19–21). To begin to distinguish between these interactions during cellular stress, we constructed eIF4G1-null (*tif4631Δ*) mutants, overexpressed *DED1* and the *ded1* mutants in these cells, and examined them for defects in growth and SG formation. *DED1* overexpression inhibited growth to a similar extent in the eIF4G1-null mutant as compared to in wild-type cells, indicating that the Ded1-eIF4G interaction is not critical for this inhibition (Figure 3A).

**Figure 3:**
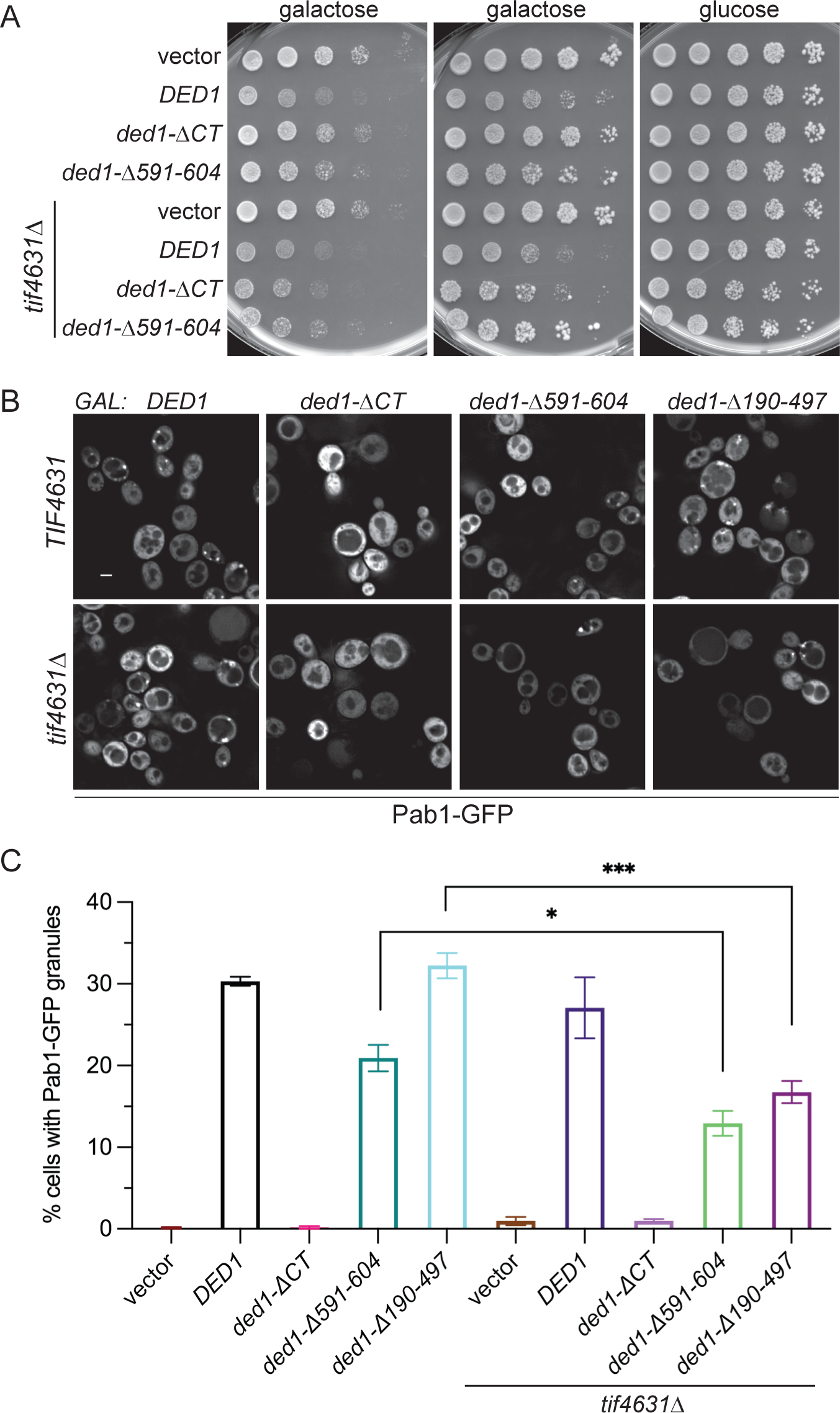
Deletion of eIF4G1 has moderate effects on GAL-DED1-induced growth inhibition and granule formation. (A) Five-fold serial dilutions of wild-type *TIF4631* (eIF4G1) or *tif4631Δ* (eIF4G1-null) cells with galactose-inducible wild-type *DED1*, *ded1-ΔCT*, or *ded1-Δ591-604* spotted on selective medium containing either galactose or glucose. Galactose plates were imaged after 2 days (*left*) or 4 days (*middle*) growth. (B) Live-cell microscopy of Pab1-GFP granules in *TIF4631* or *tif4631Δ* cells expressing galactose-inducible *DED1* or *ded1* deletion mutant constructs. Cells were grown in liquid medium containing galactose for 7 hours before imaging. Scale bar = 2 μm. (C) Quantitation of the presence of Pab1-GFP granules as the percentage of cells that contained Pab1-GFP foci. Mean and SEM of 3-7 replicates are shown. Statistical significance was determined using Student’s t-test (unpaired; * p < 0.05, *** p < 0.001).

Interestingly, however, rescue of the inhibition with expression of the *ded1-ΔCT* mutant was greatly reduced in the eIF4G1-null cells. A double mutant of *tif4631Δ* and *ded1-ΔCT* (expressed from the endogenous promoter) showed moderate synthetic growth defects even in ideal growth conditions (data not shown), so the effect of overexpressing the *ded1* mutant in eIF4G1-null cells may be due to displacing the endogenous wild-type Ded1 rather than a stress-related defect.

Examining these cells for SG formation revealed no significant difference between wild-type and eIF4G1-null cells when wild-type *DED1* is overexpressed (Figure 3B, C). Likewise, very few Pab1-GFP SGs were formed in eIF4G1-null cells when *ded1-ΔCT* was expressed, similar to wild-type cells. These results indicate that eIF4G1 is not strictly required for formation of the Ded1-induced stress granules. However, expression of the *ded1-Δ591-604* and *ded1-Δ190-497* mutants had reduced numbers of SGs in eIF4G1-null compared to in wild-type cells (Figure 3B, C). Western blotting showed roughly similar levels of Ded1, ded1-ΔCT, and ded1-Δ591-604 in wild-type and eIF4G1-null cells, arguing against effects due to protein levels for these mutants (Supplemental Figure S3). The induction of ded1-Δ190-497 protein was reduced roughly 2-fold in eIF4G1-null cells, perhaps indicating an effect of eIF4G1 on protein expression or stability, although the difference was not statistically significant. Overall, these results suggest that the mutants represent sensitized backgrounds that show that eIF4G1 indeed has an effect on SG formation, though it is moderate. Furthermore, given the differences between these mutants and *ded1-ΔCT*, it is likely that Ded1 oligomerization, the only other known C-terminal-mediated interaction, is a major contributor to SG formation.

### Ded1 promotes cell survival and growth during oxidative stress

The above approaches provided insight into Ded1-dependent mechanisms for SG formation and growth inhibition. However, overexpression of *DED1* does not recapitulate the complexity of physiological stress responses; therefore, we next examined the role of Ded1 in responding to oxidative stress through treatment of cells with hydrogen peroxide. We first tested whether Ded1 affects cell growth by measuring culture density over time in the absence and presence of peroxide (Figure 4A). In untreated cells, wild-type, *ded1-ΔCT*, and *ded1-Δ591-604* cells had similar growth rates (left panel). In peroxide-treated cultures (right panel), wild-type cells had a growth lag (λ) of about 11 hours, as calculated using the Gompertz growth equation (Supplemental Table S1A) (34). This indicated that growth was inhibited as part of the stress response, followed by an adaptation/recovery phase during which growth resumed. In contrast, the lag in *ded1-ΔCT* cells was significantly longer (22 hours) before growth resumed.

**Figure 4:**
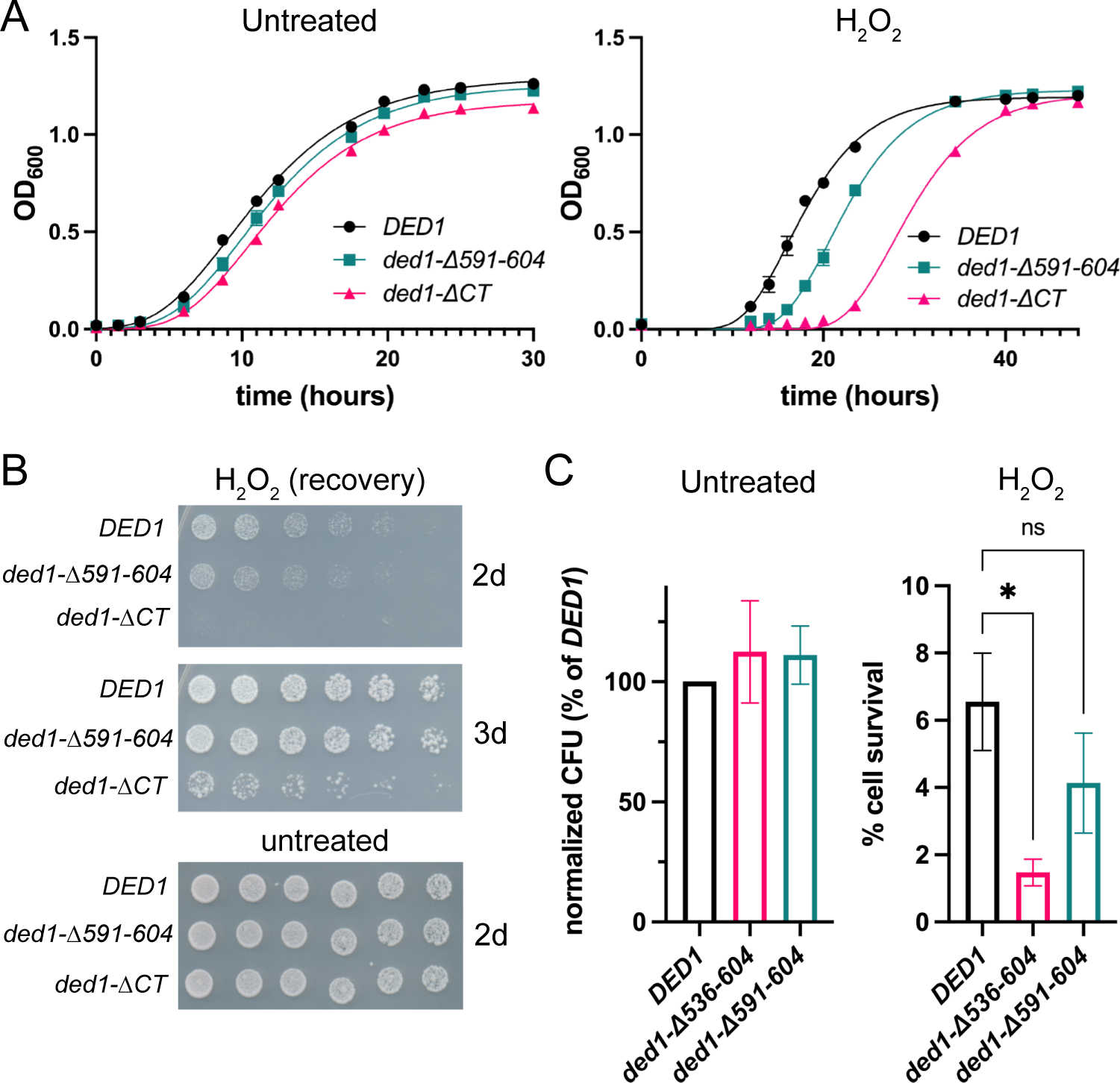
Ded1 promotes cell survival and growth during oxidative stress. (A) Growth curve analysis of *DED1, ded1-Δ591-604* and *ded1-ΔCT* strains in rich media, untreated (left) and treated with 0.8 mM H_2_O_2_ (right). Time points were fitted to the Gompertz growth equation (see Supplemental Table S1). Each time point shows the mean and SEM of 8 biological replicates performed in parallel. (B) Growth recovery of *DED1, ded1-Δ591-604* and *ded1-ΔCT* strains treated for 6 hours with 0.8 mM H_2_O_2_. A two-fold serial dilution series was performed, and cells were plated on plates lacking H_2_O_2_ and incubated for 2 or 3 days as shown. Untreated cells were diluted and plated in parallel. (C) Cell survival following 6 hours of treatment with 0.8 mM H_2_O_2_. Colony-forming units were calculated from dilutions of untreated and H_2_O_2_-treated *DED1, ded1-Δ591-604* and *ded1-ΔCT* cells after 3 days. CFUs in untreated cells are shown normalized to *DED1* to show that plating efficiency does not significantly differ between strains (left). Cell survival after treatment is shown relative to untreated CFUs for each strain (right). Data represent the mean and SEM of 4 biological replicates. Statistical significance was determined using one-way ANOVA (* p < 0.05).

Interestingly, the *ded1-Δ591-604* mutant also showed an increased growth lag compared to wild-type cells (Figure 4A, right), unlike in overexpression where it had a similar effect to wild-type *DED1*, although the defect was not as strong as in the *ded1-ΔCT* mutant. These results suggest that Ded1 and the C-terminal region have roles in cell growth during oxidative stress.

To further investigate these effects, cells were treated with peroxide, and then equal numbers of cells were plated on rich media to examine recovery of growth. As expected, peroxide treatment inhibited the growth of wild-type cells compared to untreated cells (Figure 4B, 2 days). Strikingly, peroxide-treated *ded1-ΔCT* cells displayed a severe growth inhibition compared to wild-type cells. Consistent with the growth curves, the *ded1-Δ591-604* mutant also showed moderate growth defects in this assay. These growth defects could be due either to either a delay in growth or an increase in cell death following oxidative stress. To examine the role of cell death, relative cell survival was calculated by counting colony-forming units (CFU) from wild-type and mutant strains in the presence and absence of peroxide treatment. All strains showed a significant decrease in survival after stress, but the *ded1-ΔCT* mutant had a significantly larger decrease than wild-type cells, indicating a loss of viability (Figure 4C).

However, delayed growth was eventually observed after extended incubation of the treated *ded1-ΔCT* cells (Figure 4B, 3 days), suggesting that there is also a delayed recovery of growth in these cells. Further supporting this idea, the viability of peroxide-treated *ded1-Δ591-604* cells was not significantly different from wild-type cells (Figure 4C), so the reduced growth in this mutant after stress may be largely the result of delayed recovery rather than reduced survival. Overall, these results indicate that Ded1 plays a critical role in both cell survival upon oxidative stress induction and cell growth during stress recovery.

### The role of Ded1 in stress granule dynamics during oxidative stress

Next, we examined the formation of SGs during hydrogen peroxide treatment. Peroxide treatment of wild-type cells caused an induction of SGs, defined as Pab1-GFP-positive foci, over a time course of several hours (Figure 5A, B). The percentage of cells containing SGs peaked at about 12 hours after treatment at 28%, then began to decrease, falling to near pre-treatment levels by 20 hours. Thus, the SG time-course correlates with the observed growth curve in Figure 4, with SGs increasing during the lag in growth and then decreasing as growth resumes. On the other hand, in *ded1-ΔCT* cells, SGs were more sharply induced, with over 2-fold more cells (61%) containing Pab1-GFP foci at 12 hours compared to wild-type (Figure 5B). Similar to wild-type cells, SGs began to diminish after 12 hours in *ded1-ΔCT* cells, but mutant cells continued to show a higher percentage of SGs even at later time points. Given the increased growth lag of *ded1-ΔCT* cells, this result suggests that resumption of growth correlates with eventual SG clearance. Similar to the growth defects, the *ded1-Δ591-604* mutant had slightly increased SGs compared to wild-type, although the difference was not statistically significant (Figure 5C).

**Figure 5:**
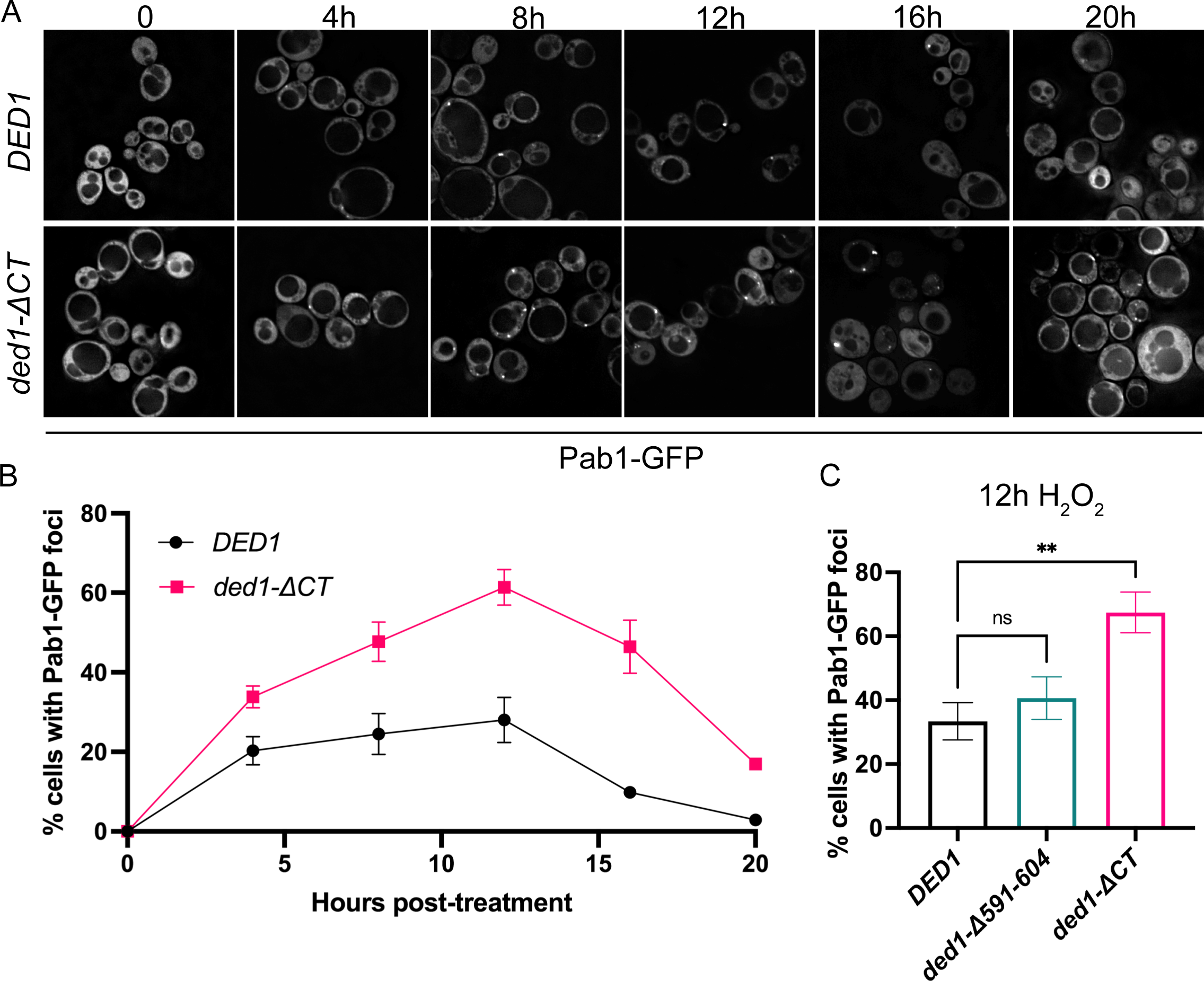
Ded1 regulates stress granule dynamics in oxidative stress. (A) Representative images of *DED1* and *ded1-ΔCT* strains expressing a Pab1-GFP reporter, after 0, 4, 8, 12, 16 and 20 hours of treatment with 0.75 mM H_2_O_2_. (B) Quantitation of Pab1-GFP foci during H_2_O_2_ time course in (A). (C) Pab1-GFP foci quantitation in *DED1, ded1-Δ591-604* and *ded1-ΔCT* strains after 12 hours of treatment with 0.75 mM H_2_O_2_. Mean and SEM of 3-4 biological replicates are shown. Statistical significance was determined using one-way ANOVA (** p < 0.01).

Notably, the *ded1-ΔCT* results in peroxide are different from the results using *ded1-ΔCT* overexpression (Figure 2). We suggest that this is a result of different means of SG induction. In contrast to the overexpression model, in oxidative stress, multiple different pathways and factors are participating in SG formation, and translation regulation can also affect SG induction (9, 35). The C-terminal region of Ded1 is thus not required for SG formation in these conditions. To examine this possibility, we expressed GFP-tagged *DED1* and *ded1-ΔCT* on low-copy plasmids in wild-type cells and treated them with peroxide. With this modest overexpression, Ded1-GFP formed some foci even in the absence of peroxide, but Ded1-positive foci increased after peroxide treatment, consistent with SG induction (Supplemental Figure S4). On the other hand, ded1-ΔCT-GFP formed multiple small speckles after peroxide treatment instead of larger discrete foci, which likely represents a defect in the ability of the mutant protein to properly condense and associate with SGs during oxidative stress.

### Ded1 interaction with eIF4G affects its role in oxidative stress

To investigate the role of Ded1’s interaction with eIF4G during oxidative stress, we examined null mutants of eIF4G1 (*tif4631Δ*) and *tif4631Δ ded1-ΔCT* double mutants for growth and SG defects. Following peroxide treatment, *tif4631Δ* cells showed a delay in growth that was intermediate (λ = 16.2h) between *ded1-ΔCT* and wild-type control cells (Figures 6A & Supplemental Table S1B), indicating that eIF4G1 also plays a role in this stress response but that the effects in the *ded1-ΔCT* mutant cannot be explained solely through Ded1 interaction with eIF4G1. Supporting this hypothesis, the *tif4631Δ ded1-ΔCT* double mutant had a similar growth delay to the *ded1-ΔCT* mutant alone (λ = 18.1h for both in this set). These results suggest an epistatic relationship between Ded1 and eIF4G wherein the effect of eIF4G on growth in peroxide is mediated through its interaction with the Ded1 C-terminal region, but the C-terminal region plays an additional role in this process, perhaps by promoting Ded1 oligomerization. The *tif4631Δ ded1-ΔCT* double mutant also showed a slight defect in the rate of growth during recovery, unlike the other mutants tested (Supplemental Table S1B). However, a similar reduced growth rate was observed in untreated mutant cells, suggesting that it is due to an effect on translation in steady-state conditions rather than stress-related (data not shown). We also examined the formation of SGs in the *tif4631Δ* mutant cells. Both *tif4631Δ* and *tif4631Δ ded1-ΔCT* mutants showed increased SGs compared to wild-type controls after 12 hours of treatment with peroxide, similar to the increase observed with *ded1-ΔCT* (Figure 6B, C). Taken together, these results suggest that the interaction of Ded1 with eIF4G1 mediates at least part of Ded1 function during oxidative stress.

**Figure 6:**
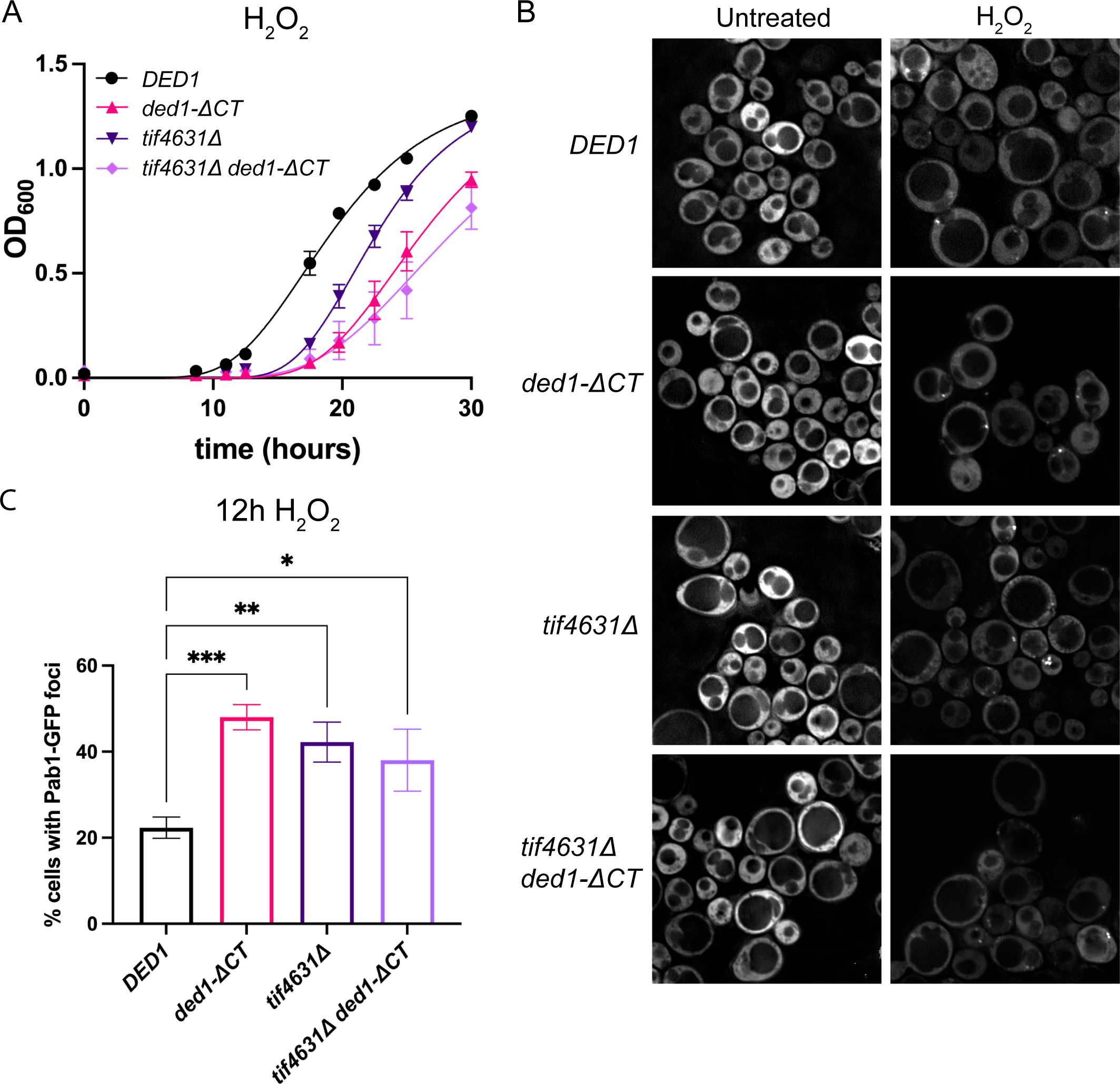
Ded1 interaction with eIF4G affects its role in oxidative stress. (A) Gompertz growth curve analysis of *DED1, ded1-ΔCT, tif4631Δ* and *tif4631Δ ded1-ΔCT* strains treated with 0.8 mM H_2_O_2_ (see also Supplemental Table S1). Each time point shows the mean and SEM of 5 biological replicates performed in parallel. (B) Representative images of *DED1, ded1-ΔCT, tif4631Δ* and *tif4631Δ ded1-ΔCT* strains expressing a Pab1-GFP reporter after 12 hours of treatment with 0.75 mM H_2_O_2_. (C) Quantitation of Pab1-GFP foci in (B). Mean of 3-4 biological replicates shown with SEM. Statistical significance was determined using a one-way ANOVA (* p < 0.05, ** p < 0.01, *** p < 0.001).

### Ded1 promotes resumption of translation during adaptation to oxidative stress

Ded1 function in translation has often focused on the dependence of mRNAs with structured 5’UTRs (16). To examine Ded1-dependent translational changes during oxidative stress, we utilized a set of previously published luciferase reporters with 5’UTRs derived from *RPL41A* that contain either a stem loop (ΔGfree = -3.7 kcal/mol) or are unstructured (Figure 7A) (24). The stem-loop containing reporter is translated less well than the unstructured one in wild-type cells, and this difference is often exacerbated in *ded1* mutants (23, 24, 36). Here, we transformed these reporters into wild-type and *ded1-ΔCT* mutant cells, treated the cells with peroxide, and performed luciferase assays over a time course. In wild-type cells, we observed significant drops in luciferase activity (to 60% of the untreated activity) with both the unstructured and structured reporters in the first two hours of treatment (Figure 7B). This reduction is consistent with the overall reduction in translation that is expected during stress conditions, although we note that the magnitude of the repression is less severe with these reporters than when bulk protein synthesis is assessed (35), likely due to the exogenous nature of the reporters. The reduction in translation in wild-type cells was maintained through 5.5 hours of treatment, but activity then increased to pre-treatment levels or above by 8 hours and was maintained through 12 hours (Figure 7B). Interestingly, this resumption of translation precedes the end of the lag phase in growth by several hours (Figure 4A & S4), suggesting that cells need this time to reshape their proteome for resumed growth. Consistent with prior studies, the structured reporter showed reduced luciferase activity compared to the unstructured one (approximately 30%) in untreated conditions (Figure 7B), but the ratio of structured to unstructured activity did not vary significantly during the time course.

**Figure 7:**
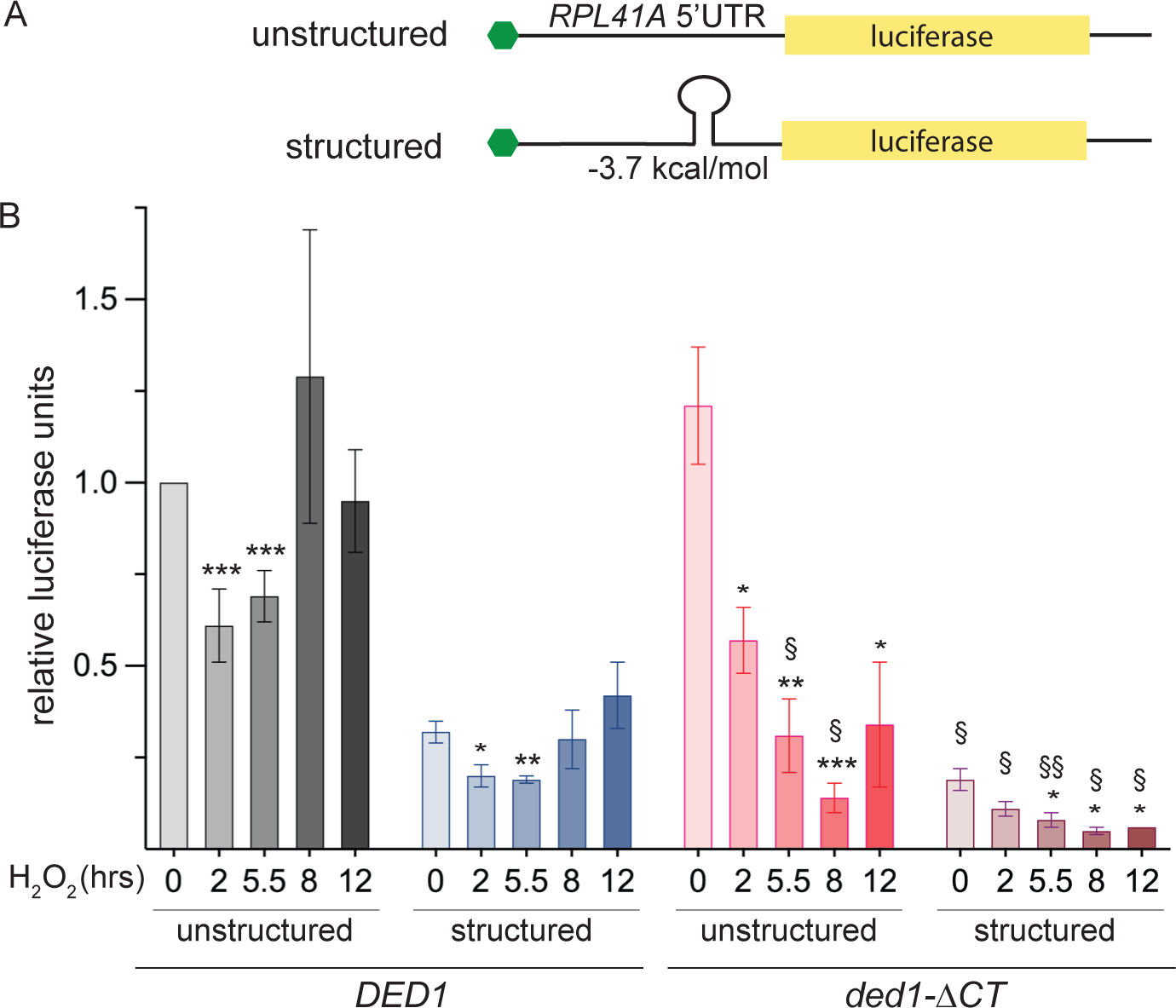
Ded1 plays multiple roles in translational regulation during oxidative stress. (A) Diagram of the unstructured (*top*) and structured (*bottom*) 5’UTR firefly luciferase reporter mRNAs. The 5’UTRs are modified versions of the yeast *RPL41A* 5’UTR; a stem-loop forming sequence is inserted in the structured reporter. (B) Time course of luciferase activity in H_2_O_2_-treated cultures of *DED1* or *ded1-ΔCT* cells containing either the unstructured or structured luciferase reporter constructs diagrammed in (A). Luciferase units obtained from each culture at each time point were normalized to the luciferase units obtained from untreated *DED1* cells containing the unstructured reporter. Mean and SEM of 3-7 biological replicates are shown. Statistical significance was determined using Student’s t-test (unpaired; * p < 0.05, ** p < 0.01, *** p < 0.001 treated vs. untreated samples from the same strain; § p < 0.05, §§ p < 0.01 *ded1-ΔCT* vs *DED1* sample).

In *ded1-ΔCT* mutant cells, activity from the unstructured reporter was similar to wild-type cells before treatment (Figure 7B). However, in treated *ded1-ΔCT* cells, translation progressively decreased over the time course through 8 hours, when translation in wild-type cells had begun to recover, and the mutant cells only partially recovered (to roughly one-third of pre-treatment levels) by 12 hours. This result is consistent with the substantially increased lag phase in the *ded1-ΔCT* cells in Figure 4, suggesting that the growth delays in the mutant may be the result of defects in resuming translation. The structured reporter in the *ded1-ΔCT* mutant showed a similar trend, although its activity did not recover to the same extent as the unstructured reporter at 12 hours (Figure 7B). Furthermore, structured reporter activity in *ded1-ΔCT* cells was significantly lower at earlier time points as well, including nearly two-fold lower in untreated cells and after 2 hours of peroxide treatment. This early reduction in translation of mRNAs with structured 5’UTRs may underlie the decreased cell survival observed in *ded1-ΔCT* mutant cells (see Figure 4C). Overall, these results suggest that Ded1 plays critical roles in regulating translation both during the initial response to stress and during the transition to resumed growth in recovering cells.

## Discussion

In this study, we examined the role of the DEAD-box RNA helicase Ded1 in stress responses using two different models. First, utilizing *DED1* overexpression, we and others observed growth inhibition, translation repression, and induction of SGs (Figure 1, 2 and (19, 26, 37)). These effects were largely mediated through the Ded1 C-terminal region, which has previously been shown to self-oligomerize and to interact with eIF4G (19, 38). Further mutant analysis indicated that the eIF4G interaction played a moderate role in growth inhibition and SG formation (Figure 3), suggesting that Ded1 oligomerization may also be a driver of these effects. Second, we subjected cells to oxidative stress through hydrogen peroxide treatment. In these conditions, Ded1 and the C-terminus were critical for cell survival and growth (Figure 4). Notably, in sharp contrast to overexpression, *ded1-ΔCT* mutant cells formed more SGs than wild-type in oxidative stress (Figure 5), suggesting different requirements for granule formation. More similar to overexpression phenotypes, both SG formation and the ability to resume growth after peroxide addition were moderately dependent on eIF4G and its interaction with Ded1 (Figure 6). Finally, in wild-type cells, translation of reporters with both structured and unstructured 5’UTRs was reduced upon oxidative stress induction but recovered after extended treatment, slightly preceding the resumption of growth, while this recovery was substantially delayed and reduced in *ded1-ΔCT* cells (Figure 7).

To integrate these disparate results, we propose the following biphasic framework for cellular stress and Ded1 function (Figure 8). The first phase is the initial response to stress conditions wherein cells shift away from pro-growth homeostasis. A number of pathways and processes are involved in this shift, resulting in inhibition of growth and proliferation as cellular resources are redirected to counter the stress (3–5). Changes in translation, specifically repression of bulk translation and upregulation of stress-specific proteins, have a critical contribution to this stress-induced growth inhibition (3, 6). SGs are also formed during this phase in response to many stresses, and SG dynamics are intricately linked to translation, e.g. translation repression is associated with SG formation and vice versa (9, 39). The second phase consists of a gradual resumption of pre-stress conditions through either removal of the stress or sufficient adaptation to it (40, 41). Cells transition from a lag phase back to growth, and general translation also resumes, after a peak in repression in the earlier response (6). Expression of stress-related proteins may also decrease, but this effect is dependent on the specific stress, its severity, and whether the stress condition is maintained. Stress granules are reduced in number during this phase (e.g. Figure 5), although to what extent this loss is due to disassembly, degradation, or simple dilution as cells begin dividing is unclear.

**Figure 8:**
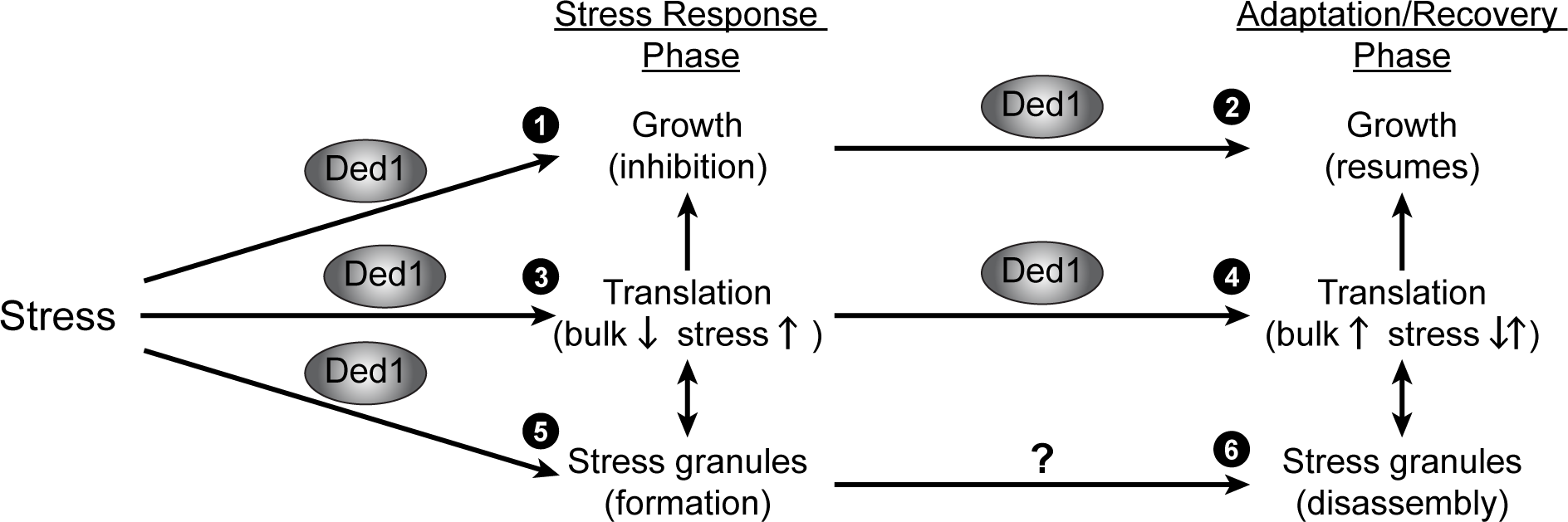
Ded1 has multiple effects during cellular stress responses. A model for Ded1 function during both the initial stress response and during adaptation/recovery. Ded1 plays roles in both translation regulation and formation of SGs during the stress response, leading to growth inhibition (1, 3, & 5). Likewise, Ded1 function is important for translation upregulation in the recovery phase, leading to resumption of growth (2, 4). A role in SG disassembly has not been identified to date (6).

Multiple lines of evidence suggest that Ded1 plays roles in several of the different aspects of stress responses described above. During the initial response phase, Ded1 functions to inhibit growth (Figure 8, #1), indicated by increased growth in *ded1* mutants (compared to wild-type) treated with rapamycin, as we showed previously (32). Likewise, the growth inhibition by *DED1* overexpression (and suppression of this phenotype in *ded1-ΔCT*) that we and others have observed is consistent with this role (Figure 1 and (19, 26, 37)). We suggest that a failure to properly halt growth during stress represents a misallocation of cellular resources, resulting in a loss of viability, as we observed in *ded1-ΔCT* mutants during both oxidative stress and nitrogen withdrawal (Figure 4C and (32)). Furthermore, the results for *ded1* mutants during oxidative stress suggest that Ded1 also plays a role in the length of the lag and the transition to resumed growth during the recovery phase (Figure 8, #2). Although *ded1-ΔCT* cells have reduced viability compared to wild-type, this mutant also has an increased lag phase and a delay in resuming growth, as observed in the growth curves and serial dilutions (Figure 4A & B). Further supporting this role, the *ded1-Δ591-604* mutant has nearly the same viability as wild-type cells but is still delayed in recovery of growth. Overall, our results suggest that Ded1 and its C-terminal region have critical roles in both growth inhibition during stress responses and in resumption of growth during recovery.

Since Ded1 is known to regulate translation, it is likely that these effects on growth by Ded1 are mediated through changes in translation. During the initial phase of the stress response, we propose that Ded1 inhibits growth through repression of general translation (Figure 8, #3).

This hypothesis has been proposed by several groups and is supported by prior results in rapamycin-treated cells, *DED1* overexpressing cells, and *in vitro* translation assays (19, 26, 32, 37). In particular, we showed previously that translation repression was reduced in *ded1-ΔCT* cells treated with rapamycin (32). Consistent with that result, here we observed defects in translation of the structured 5’UTR reporter in *ded1-ΔCT* mutant cells at all time-points during oxidative stress (Figure 7). However, while we also observed changes in translation of the unstructured reporter during the time-course, the reductions in *ded1-ΔCT* mutant cells at early-time points were similar to wild-type cells. This result may reflect limitations of the luciferase reporter to completely replicate the translational changes in the stress response; for example, the roughly two-fold reduction in translation of the reporters at early time points is much less severe than has been reported using polysome sedimentation analysis for similar peroxide concentrations (35). In any case, the severe growth inhibition of *DED1*overexpression and the substantial reduction in cell survival in *ded1-ΔCT* cells after peroxide treatment (Figures 1B & 4C) strongly suggest that Ded1 affects translation during the early portions of stress responses. In addition, given the differences between the structured and unstructured reporters, Ded1 is likely to have complex effects on different subsets of mRNAs. Guenther et al. showed that Ded1 can control the usage of alternative translation initiation sites (25), so Ded1 may have a larger repressive effect on some mRNAs and/or upregulate translation of other mRNAs, perhaps including stress-related ones.

As a second role during stress, here we also present evidence linking Ded1 to translation during the recovery phase (Figure 8, #4). In wild-type cells, translation of both the structured and unstructured reporters recovered by the 8-hour time point, preceding the end of the lag phase and resumption of growth (Figures 4 and 7). In *ded1-ΔCT* cells, however, translation progressively decreased and then recovered more slowly, consistent with the delay in growth in this mutant.

Therefore, in addition to repressing translation during the initial stress response, Ded1 (and the C-terminal region) also play a role in promoting translation during the recovery, specifically as cells transition from a stress-induced lag phase to a resumption of growth. As in the initial response (and during steady-state conditions), it is likely that Ded1 also has mRNA-specific effects during this phase. Future studies using ribosomal profiling may be able to further investigate this possibility.

Ded1 has previously been extensively linked to SGs, and it appears to play a role in their formation during stress (Figure 8, #5). This hypothesis is most directly supported by the formation of SG-like foci upon *DED1* overexpression, observed here and previously (Figure 2 and (19)), as well as a reduction in SGs following knockdown or pharmacological inhibition of its homolog DDX3X (28, 42). In addition, Ded1 and its homologs have been identified numerous times as stress granule components, including in proteomic analysis of SGs (9, 19, 28, 29, 43).

Further complementing these results, *ded1* mutations also altered SG dynamics in both the overexpression model and oxidative stress (Figures 2 & 5). However, mutants such as *ded1-ΔCT* had different effects on SG formation in the two models, decreasing SGs when overexpressed but increasing SGs upon peroxide treatment. In overexpression, SG formation is likely driven by Ded1 directly; therefore, defects in Ded1’s interactions with itself and other SG components may lead to a reduction in SGs. This effect would be minimized, however, in oxidative stress where many pathways and factors contribute to SG formation. Furthermore, changes in translation also influence SG dynamics (9, 39); thus, we suggest that the increases in SGs in *ded1* mutants treated with peroxide are caused by differences in translation in these mutants during stress.

Ded1’s specific function in SGs remains unknown and is complicated by limitations in the understanding of SGs themselves (44). As an RNA helicase, it is reasonable to speculate that Ded1 may affect the sorting of mRNAs in SGs, as recently proposed by Hondele et al. (30), but further work would be needed to examine this hypothesis. Others have suggested that Ded1 may have a role in SG disassembly during the recovery phase as well (19), but there is little supporting evidence to date. Here, we observed that SGs decrease at similar rates in wild-type and *ded1-ΔCT* cells in the late stages of oxidative stress (Figure 5B), arguing against such a role for Ded1 in SG clearance (Figure 8, #6).

The mechanism of Ded1 function in the stress response has yet to be fully defined. Here, we used different mutations in *DED1* and eIF4G in order to examine the molecular requirements for the various stress-responsive roles of Ded1. First, it is clear that the C-terminal region of Ded1 is critical to the stress response given the many defects observed in the *ded1-ΔCT* mutant. In order to begin to distinguish between the effects of eIF4G binding and Ded1 oligomerization, we utilized both deletions of eIF4G1 (*tif4631Δ*) and the *ded1-Δ591-604* mutant, which severely reduces eIF4G binding *in vitro* while having only a minor effect on oligomerization (32). In general, we observed defects in both growth and SG formation with these mutants (Figures 3 – 6), although they were more moderate than with the full C-terminal deletion mutant.

Interestingly, the *tif4631Δ ded1-ΔCT* double mutant showed similar results to the *ded1-ΔCT* mutant alone, suggesting an epistatic relationship, although interpretation of the double mutant is complicated by its growth defects even in the absence of stress. Taken together, these results indicate that eIF4G binding by Ded1 does contribute to its function in stress, which is consistent with prior studies implicating eIF4G in stress responses and in SGs (33, 45, 46). However, the more severe effects in the *ded1-ΔCT* mutant suggest that Ded1 oligomerization (or some other unknown interaction) also plays a major role, consistent with the idea that promiscuous protein-protein interactions promote SG assembly (47). Interestingly, in the overexpression model, deleting most of the central helicase domain in the *ded1-Δ190-497* mutant had similar effects to wild-type *DED1* in both growth inhibition and SG formation (Figures 1 – 3). Since this mutant is predicted to lack significant enzymatic activity and RNA binding affinity, these results suggest that neither is absolutely required for these effects, at least in the overexpression model.

However, this is unlikely to be the case for Ded1 translation regulation, and future studies will be required to further tease apart these mechanisms.

Finally, the extent to which Ded1 function in stress may be distinct in different kinds of stresses remains somewhat unclear. Several studies have now shown that Ded1 plays a role in regulating translational responses to multiple different stresses, and some aspects, such as the importance of the C-terminal region, are present in all cases to date. However, differences in Ded1 function may also be present. For example, during heat shock, a model has been proposed that as temperature rises, Ded1 condenses into SG-like structures and is not able to facilitate translation of housekeeping mRNAs with structured 5’UTRs (31). While this model is straightforward, it does not fully account for our findings here or previously in other models of stress, including the effects of *ded1* mutants on translation and SG assembly in different stresses as well as the effects of interaction with eIF4G (Figures 3, 5, 7 and (32)). As another example of the diversity of stress responses regulated by Ded1, our previous study showed that Ded1 plays a role in the response to a stress that does not induce SGs (pharmacological inhibition of TOR) (32). Furthermore, the precise subsets of mRNAs dependent on Ded1 are likely to be different in different stresses, given the need to respond to specific cellular conditions (oxidative imbalance, nutrient deprivation, heat shock, etc.). Here, the framework presented in Figure 8 is intended to be inclusive, and individual stresses might include or exclude various specific roles. Future studies will be needed to further delineate these mechanisms, particularly as the complexity of Ded1 function will likely have important implications for pathologies such as cancer and aging, where dysregulation of stress responses contributes to disease progression.

## Materials and Methods

### Yeast strains and plasmids

Yeast strains and plasmids used are listed in Supplemental Tables S2 and S3. Strains containing different *ded1* mutants under the control of the endogenous *DED1* promoter were created by plasmid shuffle starting with strain SWY4093 (*ded1::KAN +pCEN/URA3/DED1*) or strain TBY134 (*ded1::KAN, tif4631::HYG + pCEN/URA3/DED1*).

Strains that conditionally overexpressed wild-type *DED1* or *ded1* deletion mutants were created by transforming strain TBY121 (*ded1::KAN + pCEN/LEU2/ded1-ΔCT*), SWY4093, or TBY134 with plasmids that expressed *DED1* or the indicated *ded1* mutant under the control of the *GAL1/10* promoter, plus a plasmid that constitutively expressed *PAB1-GFP* where indicated.

Plasmids for galactose-inducible overexpression of untagged *DED1*, *ded1-ΔCT* or *ded1-Δ14aa* proteins *(ded1-Δ535-548*, *-Δ549-562*, *-Δ563-576*, *-Δ577-590*, and *-Δ591-604*) were constructed as follows: pTB137 was constructed by inversion of the *Xho*I fragment containing the *GAL-DED1-HHA* sequence in pRP2086 (encoding the Ded1-His-HA-ProtA fusion protein) relative to the vector backbone. Then, untagged *GAL-DED1* (pTB138), *GAL-ded1-ΔCT* (pTB148) and *GAL-ded1-Δ14aa* (pTB139 through pTB143) plasmids were constructed by replacing pTB137 sequence downstream of the internal *Bam*HI site at nt333 of the *DED1* coding sequence with the analogous sequence from plasmids pSW3619, pTB136, or pTB111 through pTB115, respectively (32, 48). Plasmid pTB144, for galactose-inducible expression of *ded1-Δ190-497*, was constructed as follows: pTB138 was digested with *Age*I (sites at nt564 and nt1488 of the *DED1* coding sequence) and the plasmid backbone plus the *DED1* N-and C-termini sequences was isolated and re-ligated, resulting in an in-frame deletion.

### Growth assays

All yeast cultures were incubated at 30°C. Serial dilution growth assays were performed as previously described (26). To analyze growth under conditions of *DED1* or *ded1* mutant overexpression, cells were pre-grown in selective SD liquid medium containing 2% sucrose, serially diluted 5-fold in SD medium, and spotted on selective SD solid medium containing 2% galactose or 2% dextrose (as control). Micro-well growth curves were generated by growing yeast cultures in YPD overnight in 96-well plates (Costar 96-well flat-bottom). For each strain 5-8 biological replicates were grown in parallel. Cells were back-diluted to OD_600nm_ of 0.1 and allowed to grow to mid-log phase. 20 μl of mid-log culture was added to 180 μl of fresh YPD with or without 0.8 mM H_2_O_2_ (Beantown Chemical, Hudson, NH) in 96-well plates. Plates were incubated at 30°C with shaking at 400 rpm to reduce cell settling. OD_600nm_ measurements were obtained at various timepoints with a VERSAmax microplate reader using SOFTmax Pro 3.1 software. For determination of growth parameters, data points were curve-fitted using Graphpad Prism to a re-parameterized form of the Gompertz growth equation (34):

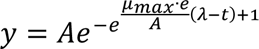

This yielded lag phase duration (λ) and maximal growth rate (μ_max_) values for each strain and condition, reported in Supplemental Table S1, and statistical significance was determined by the extra sum-of-squares F-test.

### Western blotting

For analysis of Ded1 and eIF4G1 protein levels during overexpression of *DED1* or *ded1* mutant constructs, cells were grown as described for granule analysis by fluorescence microscopy, and crude cell lysates were prepared by lysis in 1.85 M NaOH and 7.4% β-mercaptoethanol followed by trichloroacetic acid precipitation (26). Proteins were separated by SDS-PAGE, blotted, and probed with antibodies specific for Ded1 (VU318, described in (48)), eIF4G1 (gift from R. Parker (49)), Pgk1 (Life Technologies) or Pab1 (Santa Cruz). HRP-conjugated secondary antibodies were used to visualize chemiluminescent bands on a Sapphire biomolecular imager (Azure Biosystems). Ded1 band intensity was measured by densitometry using ImageJ/Fiji software and was normalized to Pgk1 band intensity in the same sample. Statistical significance was determined via Student’s t-test.

### Granule analysis by fluorescence microscopy

For stress granule analysis, the indicated strains were transformed with plasmids that expressed *PAB1-GFP* alone (pRB16) or both *PAB1-GFP* and *EDC3-mCherry* (pRP1657). For analysis of Pab1-GFP granules induced by *DED1* or *ded1* mutant overexpression, cells were cultured in selective SD media containing 2% sucrose at 30°C, back-diluted to OD_600nm_ = 0.10-0.15 in selective SD media containing 1.75% galactose + 0.25% sucrose, and cultured at 30°C for an additional 6 (TBY121-based strains) or 7 hours (SWY4093 and TBY134-based strains) prior to imaging. For analysis of Pab1-GFP granules induced by H_2_O_2_, strains were grown to OD_600nm_ = 0.2-0.4 in SD -Leu and then treated with 0.75 mM H_2_O_2_ for the indicated times. Images were captured using a DeltaVision Elite inverted microscope (Applied Precision/GE Healthcare) with an Olympus 100× plan apo NA 1.4 objective and appropriate filter sets. Z-series datasets were collected with a pco.edge sCMOS camera at a step size of 0.4 µm. Post-acquisition deconvolution was performed using SoftWorx software (Applied Precision). Z-series processing, quantitation and cropping were completed in ImageJ/Fiji; sizing and brightness adjustment were completed in Adobe Photoshop. To determine the percentage of cells with Pab1-GFP granules, the Image J/Fiji “Cell Counter” plug-in was used to track manual counts of the total number of cells, and the number of cells containing at least one Pab1-GFP focus, in merged Z-series, deconvolved images. Reported numbers for each strain undergoing either galactose induction or H_2_O_2_ treatment for the noted time represent a mean of ≥3 biological replicates. A minimum of 100 cells (mean of >300 cells) were scored for each replicate.

Statistical significance was determined by Student’s t-test or ANOVA as appropriate.

H_2_O_2_ growth/viability assays: All yeast cultures were grown to mid-log phase at 30°C in SD -Leu medium. Cultures were untreated or treated with 0.75 mM H_2_O_2_ and incubated at 30°C with shaking at 190 rpm for 6 hours. Cells were spun down and washed in SD -Leu medium and resuspended to a concentration of 1 x 10^6^ cells/ml. Two-fold serial dilutions were plated on 15cm SD -Leu agar plates. Plates were scanned after 2 and 3 days of incubation at 30°C. Cell viability was measured by counting yeast colonies in 2-fold serial dilution spots after 3 days of recovery. Fiji software was used to count colonies in serial dilution spots where individual colonies were clearly distinguishable, and colony-forming units (CFU) were calculated. The strain viability was determined by averaging normalized CFU from a minimum of 3 spots in the same dilution series for each experiment. Strain viability shown was averaged from 4 independent serial dilution experiments, and statistical significance was determined by one-way ANOVA.

### Translation scanning assays

Scanning assays for structured 5’UTR sequences were carried out similarly to (36). Briefly, cells transformed with either the control unstructured 5’UTR-firefly luciferase reporter (pFJZ342) or the stem-loop-containing 5’UTR-firefly luciferase reporter (pFJZ623) were cultured in triplicate in selective SD media at 30°C. H_2_O_2_ was added to a final concentration of 0.75 mM. Cell lysates were generated at various timepoints via bead beater in luciferase lysis buffer (25 mM Tris phosphate pH 7.8, 2 mM EGTA, 2 mM DTT, 0.5% Triton X-100, 10% glycerol). Luciferase assays were performed using a standard luciferin reagent (Promega) on a Glomax 20/20 luminometer (Promega). For each biological replicate, values obtained from the triplicate cultures were normalized to cell concentration and averaged. Statistical significance was determined by Student’s t-test.

## Acknowledgments

The authors would like to thank J. Ross Buchan, Roy Parker, Angela Hilliker, and Alan Hinnebusch for reagents, and members of the Bolger and Buchan laboratories for helpful advice and discussions. This work was supported by the National Institutes of Health (1R01-GM136827) and the American Cancer Society (RSG-1326301-RMC).

